# Variability of EEG electrode positions and their underlying brain regions: visualising gel artifacts from a simultaneous EEG-fMRI dataset

**DOI:** 10.1101/2021.03.08.434424

**Authors:** C. L. Scrivener, A. T. Reader

## Abstract

We investigated the between-subject variability of EEG electrode placement from a simultaneously recorded EEG-fMRI dataset. Neuro-navigation software was used to localise electrode positions in *xyz* and MNI space, made possible by the gel artifacts present in the structural MRI images. To assess variation in the brain regions directly underneath each electrode, we used both raw MNI coordinates and labels from the Harvard-Oxford Cortical atlas. In a sample of 20 participants, the mean standard deviation of electrode placement was 3.94 mm in *x*, 5.55 mm in *y*, and 7.17 mm in *z*, with the largest variation in parietal and occipital electrodes. In addition, the brain regions covered by electrode pairs was not always consistent; for example, the mean location of electrode P07 was mapped to BA18, whereas P08 was closer to BA19. Further, electrode C1 was mapped to the left primary motor cortex, whereas C2 was closer to right pre-motor cortex. Overall, the results emphasise the variation in electrode positioning that can be found even in a fixed cap, potentially caused by between-subject differences in brain morphology. We present a relatively simple method for approximating the location of electrodes in a simultaneous EEG-fMRI data set with accompanying analysis code, and suggest that researchers check the regions underlying their EEG ROIs to improve the generalisability and reliability of their neuroimaging results.

## 1. Introduction

Scalp electroencephalography (EEG) is one of the most frequently used neuroimaging methods, providing information about changes in electrical potential across the brain with high temporal resolution. Typical EEG setups measure activity across multiple points on the scalp. Electrodes are usually placed according to the international 10-20 system for around 21 channel recordings, 10-10 for between 64 and 85 channels, or 10-5 for high-density caps of more than 300 channels (Oostenveld et al., 2001; Jurak et al., 2007). These values refer to the distances between electrodes in relation to the total cap size (i.e., 20% of the total distance from the inion to the nasion) and aim to provide consistency across experiments. Electrodes are placed on the head of the participant with reference to anatomical landmarks such as the inion, nasion, and left and right pre-auricular points, such that the central electrode Cz is approximately aligned with the vertex. Given careful placement of the electrode cap during experimental setup, experimenters assume that the electrode placement will be roughly consistent across participants. Further, when selecting a subset of electrodes for use in EEG analysis, we assume that they are in a similar position across subjects and that we are comparing activation from similar regions of the brain.

Several studies have investigated electrode placement variations in the 10-20 (Steinmetz et al., 1989; Jack et al., 1990; Homan et al., 1997; Towle et al., 1993; Lagerlund et al., 1993; Khosla et al., 1999; Okamoto et al., 2004; Herwig et al., 2003; Atcherson et al., 2007) and 10-10 (Koessler et al., 2009) systems. For example, Okamoto et al. (2004) recorded the normalised MNI and Talairach coordinates of electrode positions across 17 participants. From the 10-20 electrode layout used, Fp1 and Fp2 had the smallest deviation of around 5 mm in their MNI coordinates (reported across the *x, y*, and *z* dimensions), compared to the largest variation of roughly 10 mm identified in occipital electrodes O1 and O2. Each electrode position was also projected onto the cortical surface to provide an estimate of the underlying brain region. Using the mean location across all participants, the electrodes largely conformed to their intended positioning; for example, P3 and P4 projected to the superior parietal lobule and precuneus, and O1 and O2 projected to the occipital gyrus and cuneus. However, the electrodes commonly used to locate the motor cortex (C3 and C4), only projected to the precentral gyrus in an average of 13% of cases. These results demonstrated the variation in location of electrodes in the 10-20 layout when collated across all participants and encourage some caution when assuming consistency in the underlying cortex.

Koessler et al. (2009) recorded the normalised Talairach coordinates of electrodes positions projected onto the cortical surface using the 10-10 electrode layout (rather than the 10-20) and therefore examined a greater number of electrodes than Okamoto et al. (2004). Across 16 participants, they reported a grand standard deviation of 4.6 mm in the *x* direction, 7.1 mm in *y*, and 7.8 mm in *z*, with variation across projected cortical positions. Fp2 had the smallest global standard deviation of 67 mm^3^ and P1 had the largest of 548 mm^3^. Some electrodes projected to the same region consistently (FP1, FP2, O1, and O2), whereas others had larger variance (C6 and FC6). For example, FP1, FP2, FC1, and FC2 projected onto the superior frontal gyrus in 100% of participants, and O1 and O2 always projected onto the occipital gyrus (BA 18: 81%, BA 19: 19%). In comparison, most central and parietal electrodes projected onto four different BA regions across participants; electrode P4 projected to BA 39 (31%), 7 (25%), 40 (25%), 19 (19%), and electrode P8 projected to BA 19 (56%), 37 (19%), 20 (12.5%), 39 (12.5%). Overall, variance in the underlying cortical regions was smallest for frontal and temporal electrodes, and greatest for central and parietal electrodes. This again suggests not only that positions vary across participants, but that the consistency of these positions is electrode and region dependent.

Whilst these results have important implications for making inferences from data derived from electrode positions, both Koessler et al. (2009) and Okamoto (2004) compared the location of manually positioned electrodes, without the aid of a cap with fixed locations. Therefore, errors in manual placement could have increased the variation in electrode location across participants. Atcherson et al. (2007) recorded the three-dimensional locations of 15 electrodes fixed within a 72 channel Neuromedical Quick Cap. Despite the addition of an electrode cap, the electrode locations had standard deviations ranging from 3 mm to 12.7 mm in pre-auricular-nasion coordinates. In this case, the largest deviations occurred in M1 and M2, placed over the mastoids, as well as FPz (the most frontal central electrode) and Iz (the most posterior occipital electrode). The largest deviations therefore occurred in the electrodes around the edge of the cap, which could be explained by variations in participant skull shapes.

Overall, several studies have provided evidence against the assumption that a chosen electrode of interest will be proximally located to the same region of cortex across participants. This is perhaps not surprising, given the potential extent of between-subject variability in the size and arrangement of the cerebral cortex. However, consistent placement of EEG electrodes is often assumed when their location is used to inform other methods. For example, the 10-20 and 10-10 electrode layouts are regularly used to guide transcranial magnetic stimulation (TMS), where stimulation sites are chosen based on the position of specific electrodes such as those over the dorsolateral prefrontal cortex (Herwig et al., 2003). Structural or functional MRI-guided TMS stimulation is often considered to be a more reliable technique (Sack et al., 2009; de Witte et al., 2018), and a recent meta-analysis of rTMS studies identified that MRI-guided targets for stimulation were associated with increased disruptive effects of TMS (Beynel et al., 2019). However, in 2016 (the latest year included in the meta-analysis), only 18% of studies used MR-guided TMS (Beynel et al., 2019). This constitutes a drop of 52% from studies between 2007 and 2013, suggesting a move back to older methods using EEG electrode guided targeting, and the need for a re-evaluation of the reliability of this method.

The aim of this study was to further understand the variability of EEG electrode positions in a commonly used research-grade EEG cap layout (BrainAmp MR, Brain Products GmbH, Gilching, Germany). We took advantage of a pre-existing neuroimaging dataset taken from a combined EEG and functional magnetic resonance imaging (fMRI) experiment, using 64 channel fixed electrode caps from Brain Products with a 10-10 electrode layout (Scrivener at al., in press). Whilst several groups have developed methods to recover EEG electrode positions from simultaneous EEG-fMRI data using specific MRI acquisition methods (Butler et al., 2018) or reconstruction from acquired structural scans (Marino et al., 2016; Silva et al., 2016; de Munck et al., 2012; Whalen et al., 2008, Koessler et al., 2008; Jurcak et al., 2005; Lamm et al., 2001; Kozinska et al., 2001; Brinkmann et al., 1998), these approaches often require methods and toolboxes that are not yet widely used. As such, we additionally provide a novel and simple way of projecting electrode locations to the cortical surface using electrode gel artifacts (that appear on the MR image underlying electrode positions) and commercially available equipment. We also provide the code to reproduce our results, or to apply to separate data sets.

This method uses a stereotactic neuro-navigation system (Brainsight, Rogue Research Inc., Montreal, QC, Canada), that has built in function to project from the scalp to the underlying cortex. Electrode gel artifacts can be visualised using the scalp reconstruction function, facilitating localisation of the electrode positions on the skull of each participant. These locations can then be projected onto the cortical surface using the inbuilt functionality of Brainsight. Using this method, we report the standard deviation of electrode positions on the skull and on the cortical surface, as well as the variability of underlying brain regions. As far as we are aware, electrode gel artifacts have not yet been used to provide a comprehensive assessment of EEG electrode position variability, either on the skull or the cortical surface, despite the fact they provide a simple method of localising brain regions under the cap.

## 2. Materials and methods

We used 20 structural scans collected for a previously reported EEG-fMRI experiment (Scrivener et al., in press), for which the data is available at https://osf.io/w6bh3/. The secondary data for the current article, as well as MATLAB scripts used to analyse the data, are freely available at https://osf.io/853kw/. Participants in the original study (Scrivener et al., in press) consented for their data to be shared anonymously, and only the defaced structural scans are freely available for download.

### 2.1. Electrode Localisation

Electrode positions were localised by author ATR using Brainsight 2.3.11 (Rogue Research Inc., Montreal, QC, Canada). The skin was reconstructed from the structural MRI scan to visualise electrode gel artifacts. Electrode positions were marked by placing targets onto the centre of the gel artifacts, orthogonal to the skin. If a gel artifact was not clearly visible, the location of the electrode was inferred based on the surrounding electrode positions (18 across all participants, and never more than five in a single participant). The positions were independently checked by author CLS, and in cases of disagreement (nine electrodes across five participants) a consensus was met.

The electrode positions were then translated onto the underlying cortical surface. To do this we projected the targets to a curvilinear brain reconstruction (created using default parameters: slice spacing = 2mm, end depth = 16mm, peel depth = 0mm) using the ‘snap to’ function. Target positions (*xyz*) on the scalp and the curvilinear brain were exported as .txt files using the Brainsight review function.

### 2.2. Data analysis

The scalp and cortical locations for each participant were translated into MNI space, using the affine transformation matrix generated by the SPM12 normalise function. This matrix provides the transformation needed to move from subject space to MNI space and allows for comparison across subjects. To plot the scalp and cortical locations, we further translated the coordinates from MNI space into *xyz* using the origin of the MNI matrix. To assess the variability of electrode positions, we calculated the mean and standard deviation of its location across participants for each electrode. This was calculated separately for scalp and cortical coordinates. Given that we had a recording of the cap size for most participants, we also extracted the locations separately for each cap size.

The brain regions at each electrode location were labelled using AtlasQuery in FSL and the Harvard-Oxford Cortical Atlas (Desikan et al., 2006; Frazier et al., 2005; Goldstein et al., 2007; Makris et al., 2006), allowing us to visualise the consistency of brain regions underlying each electrode. For each electrode in each participant, we took the highest probability region reported by the atlas. We then calculated the regions reported for each electrode across all participants as a percentage. If multiple brain regions were reported with the same (highest) probability in an electrode for a single participant, we excluded that participant for the calculation of that electrode’s underlying region. We also excluded electrodes from calculation if the atlas was not able to generate a label. Percentages were calculated based on the number of usable participants for each electrode (mean ±SD, participants = 17 ±3). We also used BioImage Suite (https://bioimagesuiteweb.github.io/webapp/) to locate the Brodmann area associated with the mean coordinates of each electrode, to supplement this information.

The scripts to reproduce these results are freely available at https://osf.io/853kw/, which can also be used on independent data. To do this, researchers should save their electrode locations into a .txt file per participant, and provide a matrix describing the transformation from subject space to MNI space (e.g., as provided by the SPM normalise function). The MATLAB script provided will extract the locations given in the .txt file, save them into a results structure, calculate summary statistics, save the results into a .csv file, and save a nifti file for each participant with the locations plotted in MNI space. An additional Bash script is provided to pass each electrode coordinate to AtlasQuery in FSL and save the output into a .txt file.

## 3. Results

### 3.1. Scalp locations

The mean electrode locations across participants can be found in Table 1. Overall, we found a grand standard deviation of 3.94 mm in *x*, 5.55 mm in *y*, and 7.17 mm in *z*. The five electrodes with the smallest overall deviation (mean SD = 4.47 mm) in *xyz* were mostly in frontal and central locations (F5, F7, FC5, FCz, FT7). The five electrodes with the largest overall deviation (mean SD = 6.78 mm) were in parietal and occipital locations (O1, P3, PO3. PO4, POz). There was no visible relationship between cap size and electrode position variability (Table 2).

**Table 1:** Mean and standard deviation of the *xyz* MNI locations for each electrode, presented separately at the scalp and on the cortex.

| Electrode | Mean (SD) |  |  |  |  |  |
| --- | --- | --- | --- | --- | --- | --- |
|  | Scalp |  |  | Cortex |  |  |
|  | x | y | z | x | y | z |
| AF3 | -31.55 (4.56) | 65.1 (4.78) | 44.6 (5.99) | -25.41 (3.99) | 54.96 (4.57) | 36.6 (5.37) |
| AF4 | 34.61 (4.76) | 65.6 (4.94) | 42.92 (7.92) | 28.06 (4.42) | 55.43 (4.28) | 35.32 (6.42) |
| AF7 | -53.89 (3.69) | 59.55 (4.31) | 8.52 (6.68) | -45.43 (4.04) | 52.57 (3.71) | 7.20 (6.01) |
| AF8 | 55.41 (4.14) | 59.76 (4.68) | 10.20 (8.88) | 46.40 (3.81) | 52.38 (4.23) | 8.78 (8.23) |
| AFZ | 0.50 (4.55) | 70.36 (4.65) | 50.59 (7.57) | 0.63 (3.84) | 57.98 (4.52) | 41.08 (6.31) |
| C1 | -31.55 (4.94) | -21.59 (7.43) | 92.04 (3.41) | -25.56 (4.77) | -23.82 (6.77) | 75.26 (2.61) |
| C2 | 28.95 (5.68) | -21.98 (7.47) | 94.55 (3.36) | 23.84 (4.52) | -24.31 (6.80) | 78.00 (3.21) |
| C3 | -59.99 (4.12) | -19.25 (7.40) | 70.68 (4.65) | -50.88 (4.53) | -21.18 (6.81) | 59.95 (3.47) |
| C4 | 58.97 (4.73) | -21.28 (6.98) | 73.84 (5.65) | 50.78 (4.61) | -23.18 (6.17) | 63.58 (4.78) |
| C5 | -76.99 (2.19) | -19.50 (6.26) | 37.19 (5.91) | -66.15 (3.20) | -20.58 (5.71) | 33.83 (5.35) |
| C6 | 77.83 (2.15) | -21.23 (5.94) | 39.99 (8.02) | 66.48 (3.11) | -21.77 (5.42) | 36.16 (6.97) |
| CP1 | -32.35 (5.32) | -51.50 (7.52) | 89.76 (3.69) | -25.64 (4.41) | -48.14 (5.73) | 71.46 (3.43) |
| CP2 | 29.61 (4.72) | -52.43 (7.98) | 92.91 (3.01) | 24.46 (3.97) | -49.71 (6.28) | 75.38 (3.28) |
| CP3 | -59.90 (4.74) | -49.84 (7.27) | 69.31 (5.29) | -49.15 (4.81) | -47.78 (5.87) | 58.42 (3.80) |
| CP4 | 56.54 (4.27) | -51.89 (7.07) | 73.89 (5.87) | 46.66 (4.34) | -48.71 (5.69) | 62.98 (4.35) |
| CP5 | -74.32 (2.58) | -49.33 (6.09) | 37.02 (8.16) | -63.88 (3.49) | -47.19 (5.69) | 33.95 (7.01) |
| CP6 | 73.44 (2.21) | -52.25 (5.56) | 41.24 (7.95) | 62.43 (3.13) | -49.07 (4.98) | 37.65 (6.65) |
| CPZ | -1.60 (4.97) | -53.38 (7.80) | 96.60 (2.90) | -0.73 (4.32) | -50.47 (6.59) | 75.85 (3.31) |
| CZ | -1.09 (4.43) | -22.15 (7.70) | 99.93 (2.66) | -0.47 (3.61) | -24.64 (6.88) | 80.16 (3.89) |
| F1 | -25.78 (4.61) | 39.37 (6.27) | 72.50 (4.61) | -20.48 (4.35) | 32.36 (5.57) | 59.68 (5.07) |
| F2 | 26.17 (4.41) | 39.90 (6.01) | 73.75 (5.62) | 20.96 (3.78) | 32.86 (5.37) | 59.93 (5.04) |
| F3 | -46.04 (3.84) | 40.62 (6.37) | 55.73 (5.58) | -38.23 (3.74) | 34.47 (5.99) | 46.92 (5.14) |
| F4 | 46.51 (4.77) | 41.51 (5.70) | 57.56 (7.15) | 38.59 (4.31) | 34.99 (5.03) | 48.23 (5.93) |
| F5 | -59.99 (2.80) | 40.30 (4.91) | 33.12 (6.10) | -50.82 (3.08) | 35.00 (4.51) | 28.66 (5.50) |
| F6 | 62.13 (3.71) | 39.34 (5.05) | 34.98 (7.61) | 52.34 (3.67) | 33.9 (4.35) | 30.57 (6.51) |
| F7 | -68.45 (2.60) | 35.32 (4.52) | 4.68 (5.56) | -56.68 (2.82) | 30.5 (4.28) | 4.33 (5.10) |
| F8 | 71.6 (3.03) | 32.35 (5.26) | 6.88 (8.00) | 59.39 (3.04) | 27.37 (4.67) | 6.97 (6.98) |
| FC1 | -29.94 (4.43) | 10.07 (7.18) | 84.65 (4.01) | -24.69 (4.32) | 5.73 (6.43) | 71.11 (3.45) |
| FC2 | 28.88 (4.51) | 9.22 (6.47) | 86.09 (4.06) | 24.09 (4.01) | 5.40 (6.14) | 72.18 (3.55) |
| FC3 | -53.84 (3.95) | 10.76 (6.85) | 65.67 (5.19) | -46.09 (4.07) | 7.35 (6.76) | 56.46 (4.20) |
| FC4 | 54.9 (4.41) | 10.08 (5.90) | 67.56 (5.81) | 47.52 (4.25) | 6.49 (5.70) | 58.34 (5.18) |
| FC5 | -71.6 (2.29) | 11.83 (5.91) | 34.94 (5.11) | -61.1 (2.74) | 8.02 (5.28) | 30.64 (5.07) |
| FC6 | 73.25 (3.46) | 9.72 (4.52) | 38.28 (8.03) | 62.59 (3.44) | 6.62 (4.52) | 33.69 (7.09) |
| FCZ | 0.02 (3.73) | 11.38 (6.8) | 91.62 (3.31) | 0.41 (3.01) | 6.77 (6.27) | 75.19 (4.06) |
| FP1 | -29.72 (5.00) | 77.81 (2.42) | 13.85 (7.37) | -24.54 (4.48) | 66.41 (2.57) | 11.97 (6.81) |
| FP2 | 29.87 (5.19) | 78.59 (3.18) | 14.59 (9.50) | 25.25 (4.43) | 66.62 (2.86) | 12.19 (7.77) |
| FPZ | -0.36 (5.19) | 83.18 (2.18) | 16.67 (8.54) | -0.29 (4.56) | 69.71 (2.39) | 13.71 (7.35) |
| FT10 | 78.78 (1.80) | 0.51 (5.48) | -32.1 (8.41) | 60.47 (5.25) | -1.67 (5.63) | -31.19 (8.88) |
| FT7 | -77.03 (2.10) | 8.63 (5.19) | 2.98 (6.14) | -63.49 (3.62) | 5.75 (4.43) | 3.02 (5.36) |
| FT8 | 79.96 (1.76) | 5.11 (4.37) | 5.06 (8.60) | 66.54 (2.85) | 2.58 (3.83) | 4.93 (8.12) |
| FT9 | -77.33 (2.65) | 1.91 (5.8) | -31.28 (6.38) | -59.09 (6.18) | 0.07 (5.75) | -30.94 (5.82) |
| FZ | 0.52 (4.56) | 43.05 (6.34) | 77.99 (5.14) | 0.88 (3.46) | 34.43 (5.64) | 62.21 (4.84) |
| O1 | -31.42 (5.64) | -109.90 (3.25) | 8.94 (11.50) | -27.11 (4.68) | -99.78 (3.60) | 6.68 (10.44) |
| O2 | 26.35 (4.88) | -110.54 (2.94) | 11.57 (11.31) | 22.51 (4.30) | -100.07 (2.74) | 8.94 (10.18) |
| OZ | -2.53 (5.50) | -114.61 (2.37) | 11.90 (11.48) | -2.49 (4.97) | -102.67 (3.09) | 9.10 (10.31) |
| P1 | -31.55 (5.73) | -77.61 (6.82) | 75.8 (6.59) | -25.9 (4.45) | -68.45 (5.63) | 61.21 (4.51) |
| P2 | 24.99 (5.99) | -78.59 (6.65) | 77.65 (6.53) | 20.8 (5.18) | -69.28 (6.73) | 64.82 (4.87) |
| P3 | -52.00 (5.32) | -77.05 (6.86) | 58.59 (8.51) | -42.75 (4.45) | -69.51 (6.00) | 49.82 (5.89) |
| P4 | 47.39 (4.70) | -78.76 (6.04) | 61.12 (7.89) | 39.28 (4.64) | -70.63 (5.65) | 52.38 (6.17) |
| P5 | -63.26 (3.79) | -77.35 (5.62) | 30.97 (9.72) | -53.84 (3.61) | -71.50 (5.22) | 28.15 (8.21) |
| P6 | 60.40 (3.56) | -78.77 (5.00) | 36.36 (9.56) | 51.14 (3.76) | -71.81 (4.69) | 32.06 (8.35) |
| P7 | -69.63 (3.32) | -73.74 (4.90) | 0.78 (10.60) | -59.17 (2.82) | -69.14 (4.46) | 0.70 (10.07) |
| P8 | 67.72 (2.35) | -75.60 (4.69) | 5.83 (9.92) | 57.52 (2.74) | -70.05 (4.18) | 4.98 (9.43) |
| PO3 | -34.84 (5.56) | -98.67 (5.19) | 41.77 (9.39) | -29.71 (4.39) | -88.53 (5.70) | 35.13 (8.07) |
| PO4 | 29.24 (5.60) | -98.85 (4.95) | 45.00 (9.87) | 25.37 (4.62) | -88.73 (5.24) | 38.16 (8.93) |
| PO7 | -54.12 (4.43) | -93.91 (3.75) | 4.79 (10.26) | -46.45 (3.26) | -86.74 (3.80) | 3.87 (9.43) |
| PO8 | 50.17 (3.84) | -95.92 (4.15) | 8.99 (10.35) | 42.78 (4.03) | -88.05 (3.60) | 7.87 (9.83) |
| POZ | -3.28 (5.69) | -101.06 (5.15) | 50.6 (9.14) | -2.76 (4.79) | -90.20 (5.71) | 42.12 (7.54) |
| PZ | -2.25 (6.16) | -80.03 (7.27) | 79.64 (6.14) | -1.94 (5.31) | -69.12 (7.17) | 66.04 (4.54) |
| T7 | -81.12 (1.60) | -20.17 (5.89) | 0.58 (7.98) | -69.72 (2.66) | -20.31 (5.20) | 0.55 (7.35) |
| T8 | 83.15 (1.08) | -23.69 (6.09) | 4.20 (9.23) | 71.04 (2.95) | -23.53 (5.69) | 3.60 (8.61) |
| TP7 | -78.17 (1.68) | -49.26 (4.7) | 0.60 (9.15) | -68.19 (2.19) | -47.06 (4.03) | 0.35 (8.25) |
| TP8 | 78.51 (1.70) | -52.05 (5.25) | 3.53 (8.75) | 67.73 (2.45) | -48.93 (5.09) | 2.77 (7.97) |
| TP9 | -73.37 (2.16) | -54.91 (4.44) | -35.64 (8.25) | -57.93 (4.26) | -52.6 (3.11) | -33.16 (7.65) |
| TP10 | 73.87 (2.55) | -57.04 (4.38) | -33.84 (10.41) | 58.93 (3.95) | -53.09 (3.59) | -31.33 (9.05) |

**Table 2:** Average standard deviation of the *xyz* MNI locations at the scalp and cortex presented separately for each cap size. Note that most participants had cap size 56, and therefore the distribution is unequal. The cap size for one participant was not recorded.

| Cap size (cm) | n | Scalp |  |  | Cortex |  |  |
| --- | --- | --- | --- | --- | --- | --- | --- |
|  |  | x | y | z | x | y | z |
| 54 | 4 | 3.89 | 3.05 | 6.21 | 3.42 | 2.85 | 5.43 |
| 56 | 12 | 4.26 | 5.54 | 7.80 | 4.26 | 5.54 | 7.00 |
| 58 | 3 | 2.42 | 5.66 | 6.02 | 3.04 | 5.12 | 5.22 |
| All | 20 | 3.94 | 5.55 | 7.17 | 3.95 | 5.09 | 6.35 |

### 3.2. Cortex locations

The mean cortical locations across participants are displayed on an MNI template brain in Figure 1, and can also be found in Table 1. Overall, we found a grand standard deviation of 3.95 mm in *x*, 5.09 mm in *y*, and 6.35 mm in *z*. The five electrodes with the smallest overall deviation (mean SD = 4.34 mm) in *xyz* were in frontal locations (F5, F7, FC5, FCz, FT7). The five electrodes with the largest overall deviation (mean SD = 6.25 mm) were in parietal and occipital areas (O1, Oz, PO3, PO4, FT10). There was no visible relationship between cap size and electrode position variability (Table 2). The cortical locations labelled using the Harvard-Oxford Cortical Atlas can be found in Table 3.

**Figure 1:**
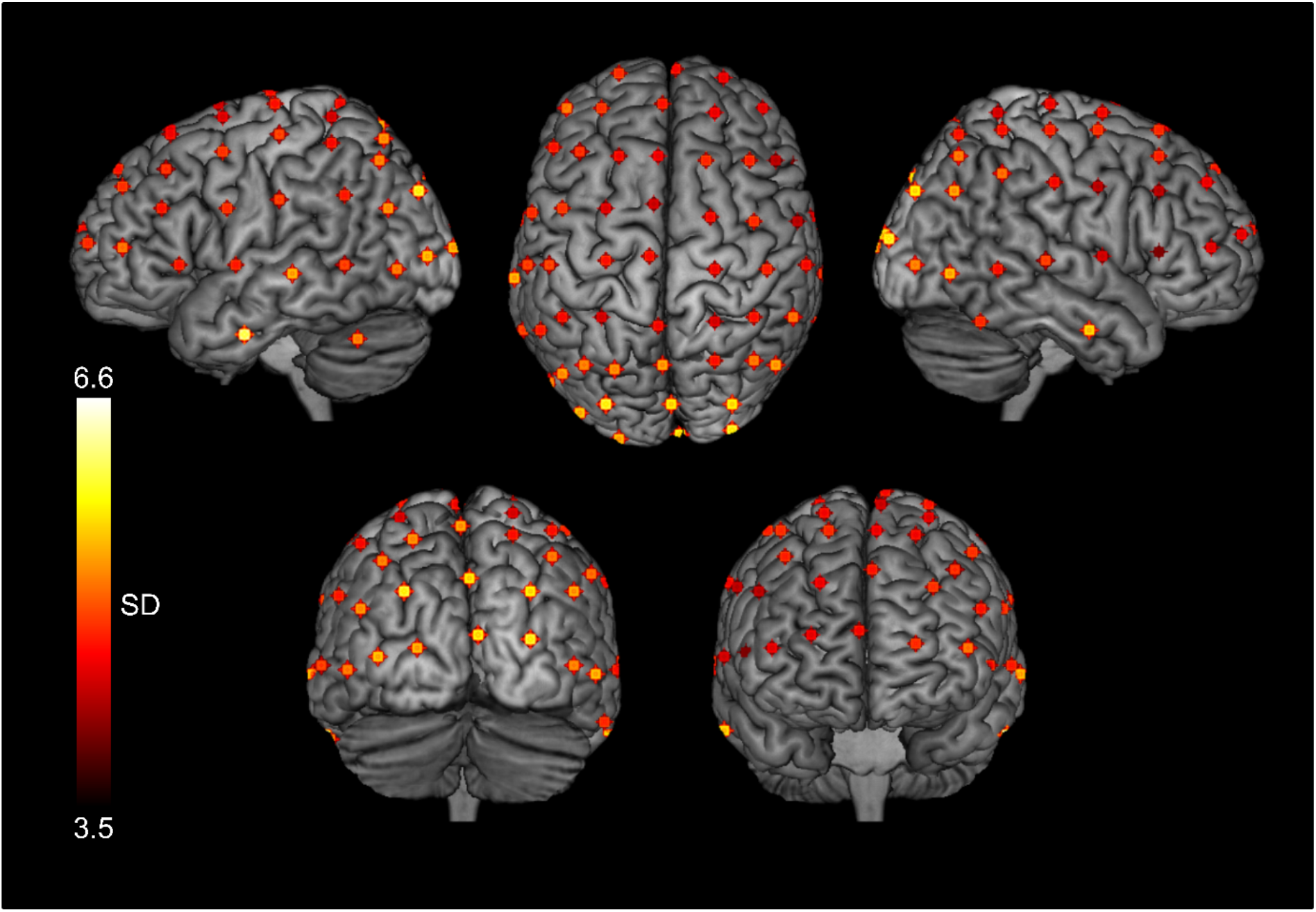
The average projected cortex locations for each of 64 electrodes across 20 subjects, displayed on an MNI template brain in MRICron. The standard deviation of each position is given by the colour, such that electrodes plotted in yellow had a higher standard deviation across subjects than those plotted in red. For visualisation purposes only, the mean co-ordinate for each electrode was convolved with a 4 mm sphere.

**Table 3:** Average electrode locations on the scalp labelled using AtlasLabel in FSL and the Harvard-Oxford Cortical Structural atlas. For each electrode we calculated the percentage of participants with each anatomical label ascribed. The closest Brodmann area for the most common cortical structure at each electrode projection is also detailed. ANG: angular gyrus, IFG: inferior frontal gyrus, ITG: inferior temporal gyrus, JLC: juxtapositional lobule cortex (formerly supplementary motor cortex), LOC: lateral occipital cortex, MFG: middle frontal gyrus, MTG: middle temporal gyrus, SFG: superior frontal gyrus, SMG: supramarginal gyrus, SPL: superior parietal lobule, STG: superior temporal gyrus

| Electrode | Percentage underlying brain regions | Brodmann area nearest mean electrode position |
| --- | --- | --- |
| AF3 | frontal pole (100%) | Left BA9 |
| AF4 | frontal pole (100%) | Right BA9 |
| AF7 | frontal pole (100%) | Left BA10 |
| AF8 | frontal pole (100%) | Right BA10 |
| AFZ | frontal pole (62.5%), SFG (37.5%) | Left BA9 |
| C1 | precentral gyrus (71%), postcentral gyrus (29%) | Left BA4 |
| C2 | precentral gyrus (87.5%), postcentral gyrus (12.5%) | Right BA6 |
| C3 | postcentral gyrus (87.5%), precentral gyrus (12.5%) | Left BA1 |
| C4 | postcentral gyrus (85%), precentral gyrus (15%) | Right BA1 |
| C5 | postcentral gyrus (50%), anterior SMG (44.4%), precentral gyrus (5.6%) | Left BA40 |
| C6 | anterior SMG (72%), postcentral gyrus (28%) | Right BA40 |
| CP1 | SPL (66.67%), postcentral gyrus (27.78%), superior LOC (5.56%) | Left BA7 |
| CP2 | SPL (80%), postcentral gyrus (13%), superior LOC (7%) | Right BA7 |
| CP3 | posterior SMG (47%), SPL (27%), ANG (13%), postcentral gyrus (13%) | Left BA40 |
| CP4 | ANG (50%), SPL (42%), postcentral gyrus (8%) | Right BA7 |
| CP5 | posterior SMG (61.11%), anterior SMG (16.67%), ANG (11.11%),<br>posterior STG (5.56%), superior LOC (5.56%) | Left BA39 |
| CP6 | ANG (53%), posterior SMG (40%), superior LOC (7%) | Right BA39 |
| CPZ | postcentral gyrus (62.5%), precuneous cortex (18.75%), SPL (12.5%),<br>superior LOC (6.25%) | Left BA7 |
| CZ | precentral gyrus (87%), postcentral gyrus (13%) | Right BA4 |
| F1 | SFG (71%), frontal pole (29%) | Left BA6/BA8 |
| F2 | SFG (77%), frontal pole (23%) | Right BA6/BA8 |
| F3 | MFG (64%), frontal pole (36%) | Left BA9 |
| F4 | frontal pole (53%), MFG (47%) | Right BA9 |
| F5 | MFG (65%), frontal pole (35%) | Left BA9 |
| F6 | MFG (50%), frontal pole (50%) | Right BA9 |
| F7 | IFG (pars triangularis) (83%), frontal pole (11%), IFG (pars opercularis) (6%) | Left BA45 |
| F8 | IFG (pars triangularis) (66.67%), IFG (pars opercularis) (16.67%), MFG (5.56%), precentral gyrus (5.56%), frontal pole (5.56%) | Right BA45 |
| FC1 | SFG (93.75%), MFG (6.25%) | Left BA6 |
| FC2 | SFG (82%), precentral gyrus (18%) | Right BA6 |
| FC3 | MFG (89%), precentral gyrus (11%) | Left BA6 |
| FC4 | MFG (92%), precentral gyrus (8%) | Right BA6 |
| FC5 | precentral gyrus (88%), MFG (6%), IFG (pars opercularis) (6%) | Left BA6 |
| FC6 | precentral gyrus (81%), postcentral gyrus (13%), MFG (6%) | Right BA6 |
| FCZ | JLC (66.7%), SFG (33.3%) | Left BA6 |
| FP1 | frontal pole (100%) | Left BA10 |
| FP2 | frontal pole (100%) | Right BA10 |
| FPZ | frontal pole (100%) | Left BA10 |
| FT10 | anterior MTG (66.7%), posterior ITG (11.1%), temporal pole (11.1%), posterior MTG (5.6%), anterior ITG (5.6%) | Right BA21 |
| FT7 | precentral gyrus (64.29%), temporal pole (14.29%), anterior STG (14.29%), IFG (pars opercularis) (7.14%) | Left BA44 |
| FT8 | precentral gyrus (62.5%), anterior STG (25%), central opercular cortex (6.25%), posterior MTG (6.25%) | Right BA6 |
| FT9 | anterior MTG (42%), temporal pole (37%), posterior MTG (16%), anterior ITG (5%) | Left BA38 |
| FZ | SFG (100%) | Left BA6 |
| O1 | occipital pole (100%) | Left BA18 |
| O2 | occipital pole (100%) | Right BA18 |
| OZ | occipital pole (100%) | Left BA18 |
| P1 | superior LOC (95%), SPL (5%) | Left BA7 |
| P2 | superior LOC (100%) | Right BA7 |
| P3 | superior LOC (95%), ANG (5%) | Left BA39 |
| P4 | superior LOC (100%) | Right BA39 |
| P5 | superior LOC (89.47%), inferior LOC (5.26%), ANG (5.26%) | Left BA39 |
| P6 | superior LOC (100%) | Right BA39 |
| P7 | inferior LOC (75%), superior LOC (20%), MTG (temporooccipital part) (5%) | Left BA19 |
| P8 | inferior LOC (85%), superior LOC (10%), ANG (5%) | Right BA19 |
| PO3 | superior LOC (65%), occipital pole (35%) | Left BA19 |
| PO4 | occipital pole (58%), superior LOC (42%) | Right BA19 |
| PO7 | inferior LOC (80%), superior LOC (15%), occipital pole (5%) | Left BA18 |
| PO8 | occipital pole (33.33%), superior LOC (33.33%), inferior LOC (33.33%) | Right BA19 |
| POZ | cuneal cortex (37.5%), occipital pole (31.3%), superior LOC (25%),<br>precuneous cortex (6.3%) | Left BA19 |
| PZ | precuneous cortex (60%), superior LOC (33.33%), SPL (6.66%) | Left BA7 |
| T7 | posterior STG (72.2%), posterior MTG (16.6%), anterior STG (5.6%),<br>anterior SMG (5.6%) | Left BA21 |
| T8 | posterior STG (52.9%), posterior MTG (29.4%), planum temporale<br>(5.9%), anterior SMG (5.9%), central opercular cortex (5.9%) | Right BA22 |
| TP7 | MTG (temporooccipital part) (73.7%), posterior SMG (15.8%), posterior<br>MTG (10.5%) | Left BA21 |
| TP8 | MTG (temporooccipital part) (75%), ANG (20%), posterior SMG (5%) | Right BA37 |
| TP9 | ITG (temporooccipital part) (100%) | Not applicable (cerebellum) |
| TP10 | ITG (temporooccipital part) (89%), MTG (temporooccipital part) (11%) | Not applicable (cerebellum) |

## 4. Discussion

We evaluated the variability of EEG electrode positions and their underlying brain regions using data recorded during a simultaneous EEG-fMRI experiment with Brain Products MR 64 channel caps. Overall, we found variance in electrode placement that was comparable with previous studies, with the largest deviations in the *z* dimension and in occipital and parietal electrodes. Consistent with previous findings, frontal electrodes had the smallest deviation across subjects, in co-ordinates both at the scalp and projected onto the brain (Okamoto et al., 2004; Koessler et al., 2009). However, we did not identify any greater variation specifically in electrodes around the edge of the electrode cap, as previously found (Atcherson et al., 2007). We also did not find any consistent effect of cap size. However, as most participants required the average cap size of 56, there were few data points from which to draw conclusions. In the future a more thorough examination of the influence of cap size on electrode position variability would be beneficial. In addition, we present a relatively simple method for approximating the location of electrodes using electrode gel artifacts, and provide the necessary analysis code for comparing scalp and cortex locations across subjects.

These results have particularly important implications for studies using TMS. It is generally proposed that MRI-guided stimulation is the most reliable approach to TMS (Sack et al., 2009; de Witte et al., 2018; Bergmann & Hartwigsen, 2020), and it is associated with increased disruptive effects (Beynel et al., 2019). However, it remains common practice to use the international 10-10 and 10-20 layout systems to guide positioning for TMS stimulation, particularly when neuro-navigation using structural or functional MRI scans is not possible (Beynel et al., 2019). This provides an approximate estimation of ROIs without the need for expensive MRI scanning time and will therefore be necessary for some experiments. Our results suggest that using EEG electrode position guided TMS may be more reliable for frontal electrodes, given the relatively small standard deviation found across participants. However, large variation in the electrode position and underlying brain regions were found for electrodes at the back of the head, including occipital and parietal ROIs, which may lead to larger between-subject differences in cortex stimulation with TMS.

Researchers also use the 10-20 layout to inform electrode choice in EEG analysis. In accordance with previous results (Okamoto et al., 2004), electrode pairs C1/C2 and C3/C4 were not reliable for approximating the location of the motor cortex across subjects. The mean locations of C3 and C4 were closer to the post-central gyrus, and while neighbouring electrode C1 was proximally located to the motor cortex, its pair electrode C2 was closer to the pre-motor cortex. Similarly, the mean location of electrode PO7 was mapped to BA18, whereas PO8 was closer to BA19. In this case, it may be beneficial to select the most relevant electrodes on an individual participant basis to calculate power or evoked potentials arising from the primary visual cortex, rather than selecting PO7 and PO8 by default. Furthermore, source localisation of EEG data is frequently used to provide an estimate of where in the brain a given change in electrical potential arises. However, interpreting source localisation at the group level could be limited by the assumption that the relationship between electrode position and underlying cortical tissue is consistent across individuals (Dalal et al., 2014; Milan et al., 2018). Of course, electrical activity recorded at the level of the scalp is the summation of activity from multiple sources on the underlying cortex, and is not exclusively representing the neural activity in the closest region of the cortex (Nunez & Srinivasan, 2006). However, researchers generally select electrodes for analysis based on their proximity to the brain region of interest.

In addition to providing the results for one EEG-fMRI data set, we highlight a user-friendly way of using electrode gel artifacts to localise electrode positions across participants. This method takes advantage of existing functions in Brainsight; a software commonly used for neuro-navigation in TMS, and therefore accessible for many neuroimaging centres. Although it is time consuming to manually label the position of each electrode for each participant, researchers could instead label a subset of electrodes for analysis (if not all are used). In this case, electrode positions were labelled after completion of the experiment. However, researchers can use the functionality of Brainsight to mark the position of some/all electrodes on the EEG cap of each participant before beginning their experiment.

As this method requires manual marking of electrode positions on the reconstructed scalp of the participant, error can be introduced by the subjective decision of the researcher. To combat this, every electrode position was checked and agreed on by both authors. A total of nine electrodes across five participants were re-labelled during this checking procedure, all of which were more difficult to visualise given a very small or very large gel artifact. However, most positions were clearly visible on the Brainsight reconstruction, and the researchers agreed on the target locations of most electrodes. An additional source of variance could arise from the choice of atlas used for analysis. We used the Harvard-Oxford cortical atlas and Brodmann regions to label the cortex underlying each electrode. The choice of atlas will influence the exact labelling, and we therefore chose a commonly used atlas available in FSL. Other researcher may choose to use atlases frequently used in their area of research, or those which detail their specific region of interest.

Overall, our results emphasise the variation in electrode positioning that can be found even using a fixed EEG cap, most likely caused by between-subject differences in brain morphology. These results are likely to vary across experiment and participant group, however, we provide an example case to demonstrate the potential variation in electrode positioning and underlying cortex across a sample group. We present a relatively simple method for approximating the location of electrodes in a simultaneous EEG-fMRI dataset with accompanying analysis code, and suggest that researchers check the regions underlying their EEG ROIs to improve the generalisability and reliability of their results.

## References

Atcherson, S. R., Gould, H. J., Pousson, M. A., & Prout, T. M. (2007). Variability of Electrode Positions Using Electrode Caps. Brain Topography, 20(2), 105–111. https://doi.org/10.1007/s10548-007-0036-z

Beynel, L., Appelbaum, L. G., Luber, B., Crowell, C. A., Hilbig, S. A., Lim, W., Nguyen, D., Chrapliwy, N. A., Davis, S. W., Cabeza, R., Lisanby, S. H., & Deng, Z.-D. (2019). Effects of online repetitive transcranial magnetic stimulation (rTMS) on cognitive processing: A meta-analysis and recommendations for future studies. Neuroscience & Biobehavioral Reviews, 107, 47–58. https://doi.org/10.1016/j.neubiorev.2019.08.018

Brinkmann, B. H., O’Brien, T. J., Dresner, M. A., Lagerlund, T. D., Sharbrough, F. W., & Robb, R. A. (1998). Scalp-Recorded EEG Localization in MRI Volume Data. Brain Topography, 10(4), 245–253. https://doi.org/10.1023/A:1022266822252

Butler, R., Gilbert, G., Descoteaux, M., Bernier, P.-M., & Whittingstall, K. (2017). Application of polymer sensitive MRI sequence to localization of EEG electrodes. Journal of Neuroscience Methods, 278, 36–45. https://doi.org/10.1016/j.jneumeth.2016.12.013

Dalal, S. S., Rampp, S., Willomitzer, F., & Ettl, S. (2014). Consequences of EEG electrode position error on ultimate beamformer source reconstruction performance. Frontiers in Neuroscience, 8. https://doi.org/10.3389/fnins.2014.00042

de Munck, J. C., van Houdt, P. J., Verdaasdonk, R. M., & Ossenblok, P. P. W. (2012). A semi-automatic method to determine electrode positions and labels from gel artifacts in EEG/fMRI-studies. NeuroImage, 59(1), 399–403. https://doi.org/10.1016/j.neuroimage.2011.07.021

De Witte, S., Klooster, D., Dedoncker, J., Duprat, R., Remue, J., & Baeken, C. (2018). Left prefrontal neuronavigated electrode localization in tDCS: 10–20 EEG system versus MRI-guided neuronavigation. Psychiatry Research: Neuroimaging, 274, 1–6. https://doi.org/10.1016/j.pscychresns.2018.02.001

Desikan, R. S., Ségonne, F., Fischl, B., Quinn, B. T., Dickerson, B. C., Blacker, D., Buckner, R. L., Dale, A. M., Maguire, R. P., Hyman, B. T., Albert, M. S., & Killiany, R. J. (2006). An automated labeling system for subdividing the human cerebral cortex on MRI scans into gyral based regions of interest. NeuroImage, 31(3), 968–980. https://doi.org/10.1016/j.neuroimage.2006.01.021

Frazier, J. A., Chiu, S., Breeze, J. L., Makris, N., Lange, N., Kennedy, D. N., Herbert, M. R., Bent, E. K., Koneru, V. K., Dieterich, M. E., Hodge, S. M., Rauch, S. L., Grant, P. E., Cohen, B. M., Seidman, L. J., Caviness, V. S., & Biederman, J. (2005). Structural Brain Magnetic Resonance Imaging of Limbic and Thalamic Volumes in Pediatric Bipolar Disorder. American Journal of Psychiatry, 162(7), 1256–1265. https://doi.org/10.1176/appi.ajp.162.7.1256

Goldstein, J. M., Seidman, L. J., Makris, N., Ahern, T., O’Brien, L. M., Caviness, V. S., Kennedy, D. N., Faraone, S. V., & Tsuang, M. T. (2007). Hypothalamic Abnormalities in Schizophrenia: Sex Effects and Genetic Vulnerability. Biological Psychiatry, 61(8), 935–945. https://doi.org/10.1016/j.biopsych.2006.06.027

Herwig, U., Satrapi, P., & Schönfeldt-Lecuona, C. (2003). Using the International 10-20 EEG System for Positioning of Transcranial Magnetic Stimulation. Brain Topography, 16(2), 95–99. https://doi.org/10.1023/B:BRAT.0000006333.93597.9d

Khosla, D., Don, M., & Kwong, B. (1999). Spatial mislocalization of EEG electrodes –effects on accuracy of dipole estimation. Clinical Neurophysiology, 110(2), 261–271. https://doi.org/10.1016/S0013-4694(98)00121-7

Koessler, L., Benhadid, A., Maillard, L., Vignal, J. P., Felblinger, J., Vespignani, H., & Braun, M. (2008). Automatic localization and labeling of EEG sensors (ALLES) in MRI volume. NeuroImage, 41(3), 914–923. https://doi.org/10.1016/j.neuroimage.2008.02.039

Koessler, L., Maillard, L., Benhadid, A., Vignal, J. P., Felblinger, J., Vespignani, H., & Braun, M. (2009). Automated cortical projection of EEG sensors: Anatomical correlation via the international 10–10 system. NeuroImage, 46(1), 64–72. https://doi.org/10.1016/j.neuroimage.2009.02.006

Lamm, C., Windischberger, C., Leodolter, U., Moser, E., & Bauer, H. (2001). Co-Registration of EEG and MRI Data Using Matching of Spline Interpolated and MRI-Segmented Reconstructions of the Scalp Surface. Brain Topography, 14(2), 93–100. https://doi.org/10.1023/A:1012988728672

Makris, N., Goldstein, J. M., Kennedy, D., Hodge, S. M., Caviness, V. S., Faraone, S. V., Tsuang, M. T., & Seidman, L. J. (2006). Decreased volume of left and total anterior insular lobule in schizophrenia. Schizophrenia Research, 83(2), 155–171. https://doi.org/10.1016/j.schres.2005.11.020

Marino, M., Liu, Q., Brem, S., Wenderoth, N., & Mantini, D. (2016). Automated detection and labeling of high-density EEG electrodes from structural MR images. Journal of Neural Engineering, 13(5), 056003. https://doi.org/10.1088/1741-2560/13/5/056003

Nunez, P. L., & Srinivasan, R. (2006). Electric Fields of the Brain: The Neurophysics of EEG. Oxford University Press.

Okamoto, M., Dan, H., Sakamoto, K., Takeo, K., Shimizu, K., Kohno, S., Oda, I., Isobe, S., Suzuki, T., Kohyama, K., & Dan, I. (2004). Three-dimensional probabilistic anatomical cranio-cerebral correlation via the international 10–20 system oriented for transcranial functional brain mapping. NeuroImage, 21(1), 99–111. https://doi.org/10.1016/j.neuroimage.2003.08.026

Oostenveld, R., & Praamstra, P. (2001). The five percent electrode system for high-resolution EEG and ERP measurements. Clinical Neurophysiology, 112(4), 713–719. https://doi.org/10.1016/S1388-2457(00)00527-7

Sack, A. T., Cohen Kadosh, R., Schuhmann, T., Moerel, M., Walsh, V., & Goebel, R. (2009). Optimizing Functional Accuracy of TMS in Cognitive Studies: A Comparison of Methods. Journal of Cognitive Neuroscience, 21(2), 207–221. https://doi.org/10.1162/jocn.2009.21126

Scrivener, C. L., Malik, A., Lindner, M., & Roesch, E. B. (2020). Sensing and seeing associated with overlapping occipitoparietal activation in simultaneous EEG-fMRI. https://doi.org/10.1101/2020.07.08.193326

Taberna, G. A., Marino, M., Ganzetti, M., & Mantini, D. (2019). Spatial localization of EEG electrodes using 3D scanning. Journal of Neural Engineering, 16(2), 026020. https://doi.org/10.1088/1741-2552/aafdd1

Whalen, C., Maclin, E. L., Fabiani, M., & Gratton, G. (2008). Validation of a method for coregistering scalp recording locations with 3D structural MR images. Human Brain Mapping, 29(11), 1288–1301. https://doi.org/10.1002/hbm.20465

